# The Impact of Spatial Normalization for Functional Magnetic Resonance Imaging Data Analyses Revisited

**DOI:** 10.1101/272302

**Authors:** Jason F. Smith, Juyoen Hur, Claire M. Kaplan, Alexander J. Shackman

**Author notes:** **Address Correspondence to:** Jason F. Smith or Alexander J. Shackman, Biology-Psychology Building, University of Maryland, College Park MD 20742 USA.

## Abstract

Spatial normalization—the process of aligning anatomical or functional data acquired from different individuals to a common stereotaxic atlas—is routinely used in the vast majority of functional neuroimaging studies, with important consequences for scientific inference and reproducibility. Although several approaches exist, multi-step techniques that leverage the superior contrast and spatial resolution afforded by T1-weighted anatomical images to normalize echo planar imaging (EPI) functional data acquired from the same individuals (T1EPI) is now standard. Yet, recent work suggests that direct alignment of functional data to a T2*-weighted template without recourse to an anatomical image—an EPI only (EPIO) approach—enhances normalization precision. This counterintuitive claim is intriguing, suggesting that a change in standard practices may be warranted. Here, we re-visit these conclusions, extending prior work to encompass newly developed measures of normalization precision, accuracy, and ‘real-world’ statistical performance for the standard EPIO and T1EPI pipelines implemented in SPM12, a recently developed variant of the EPIO pipeline, and a novel T1EPI pipeline incorporating ‘best practice’ tools from multiple software packages. The multi-tool T1EPI pipeline was consistently the most precise, most accurate, and resulted in the largest *t* values at the group level, in some cases dramatically so. The three SPM-based pipelines exhibited more modest and variable differences in performance relative to each another, with the widely used T1EPI pipeline showing the second best overall precision and accuracy, and the recently developed EPIO pipeline generally showing the poorest overall performance. The results demonstrate that standard pipelines can be easily improved and we encourage researchers to invest the resources necessary to do so. The multi-tool pipeline presented here provides a framework for doing so. In addition, the novel performance metrics described here should prove useful for reporting and validating future methods for pre-processing functional neuroimaging data.

## INTRODUCTION

Spatial normalization—the process of aligning data acquired from individual participants to a common stereotaxic atlas^1^—is routinely used in the vast majority of neuroimaging studies. Spatial normalization can be conceptualized as having two, partially overlapping aims: *precision*, indexed by the degree of agreement among individual participants’ data, and *accuracy*, indexed by the degree of agreement between the participants and the chosen neuroanatomical atlas. Both of these aims are important to consider when evaluating spatial normalization quality.

Maximizing normalization *precision* requires minimizing the degree of residual spatial variability across participants. Inter-participant spatial overlap enables voxelwise statistical analyses across participants (i.e., ‘group’ or ‘second-level’ analyses); if individual participant functional images and/or the statistical maps calculated from them are in voxelwise agreement, then to a first approximation, a given voxel (i.e., stereotaxic coordinate) in a given participant’s brain can be considered analogous to that same voxel in another’s (Fox, 1995; Fox, Mintun, Reiman, & Raichle, 1988; but see Stelzer, Lohmann, Mueller, Buschmann, & Turner, 2014; Turner, 2016; Turner & Geyer, 2014 for important caveats). This allows statistical parametric maps to be combined across participants and permits voxelwise population-level inferences (Friston et al., 1990; Friston, Frith, Turner, & Frackowiak, 1995; Holmes & Friston, 1998; Worsley & Friston, 1995). In addition to algorithmic advances, work aimed at improving the precision of spatial normalization have has focused on developing specialized target templates for particular populations (e.g., ethnic groups or youth) or study-specific samples (e.g., Buckner et al., 2004; Burgund et al., 2002; Fonov et al., 2011; Huang et al., 2010).

In contrast, the goal of normalization *accuracy* is to standardize spatial locations across studies. Maximizing accuracy requires minimizing the degree of residual variability between the functional data at hand and the target atlas. High normalization accuracy enables the use of a wide range of tools for sharing, interrogating, and interpreting neuroimaging data, including reporting locations in standardized stereotaxic coordinates as well as using pre-defined anatomical labels, probabilistic anatomical regions, tissue prior probability maps, and coordinate-based meta-analytic information (e.g., Desikan et al., 2006; Destrieux, Fischl, Dale, & Halgren, 2010; Fox, Lancaster, Laird, & Eickhoff, 2014; Laird, Lancaster, & Fox, 2005; Pauli, Nili, & Tyszka, 2017; Theiss, Ridgewell, McHugo, Heckers, & Blackford, 2017; Tyszka & Pauli, 2016; Yarkoni, Poldrack, Nichols, Van Essen, & Wager, 2011). Naturally, these approaches are only valid to the extent that the data are adequately aligned to the atlas (e.g., Hrybouski et al., 2016; Tillman et al., 2017). A normalization method can be highly accurate—with the centroid of a group of normalized participants perfectly matching the chosen atlas—yet highly imprecise, such that no individual participant matches another. Likewise, a normalization method can be highly precise—such that all of the participants perfectly aligned to one another—in a location that bears little relation to the atlas.

Given the fundamental importance of spatial normalization, a number of approaches and tools have been developed and evaluated over the past two decades (e.g., Ardekani et al., 2005; Chakravarty et al., 2009; Fu et al., 2017; Hellier et al., 2003; Klein et al., 2009; Klein et al., 2010; Yassa & Stark, 2009). A simple, but no longer common approach for functional MRI (fMRI) data is to directly align the individual T2*-weighted echo planar images (EPI) to a T2*-weighted template (e.g., Friston, Ashburner, et al., 1995; Woods, Cherry, & Mazziotta, 1992)^2^. This ‘EPI only’ (EPIO) approach has the virtue of simplicity—it requires only the fMRI data and a single optimization. Moreover, if the target template is undistorted, EPIO in conjunction with nonlinear spatial transformations has the potential to correct geometric distortions caused by inhomogeneity of the static field or regional differences in magnetic susceptibility (e.g., Chambers, Bhushan, Haldar, Leahy, & Shattuck, 2015; Wang et al., 2017). However, the location and magnitude of distortions and susceptibility-induced signal loss are known to vary with imaging parameters and across individuals (Chen, Dickey, Yoo, Guttmann, & Panych, 2003; Olman, Davachi, & Inati, 2009; Robinson, Windischberger, Rauscher, & Moser, 2004). If the distortions and signal loss embodied in the EPI template are dissimilar from those characterizing a particular dataset, ‘aggressive’ nonlinear transformations can erroneously stretch signals from adjacent tissue to ‘fill in’ regions of signal loss and reduce accuracy. More generally, the limited tissue contrast and coarse spatial resolution (~2-4 mm^3^) of typical T2*-weighted EPI sequences ultimately limit the precision of EPIO approaches.

Consequently, most investigators now rely on multistep approaches that leverage the superior tissue contrast and spatial resolution (~0.5-l mm^3^) afforded by T1-weighted ‘anatomical’ images to spatially normalize EPI ‘functional’ data acquired from the same individual, or what we term the ‘T1EPI’ approach. Here, the T1-weighted and EPI data are first co-registered. Next, an affine and/or nonlinear spatial transformation is computed for aligning the T1-weighted image to a T1-weighted template. Finally, this transformation is applied to the co-registered EPI data (Seitz et al., 1990; Woods, Grafton, Watson, Sicotte, & Mazziotta, 1998). While necessarily more complex than EPIO, T1EPI has the advantage of utilizing images with enhanced contrast and resolution as well as reduced distortion to normalize the individual functional datasets.

The co-registration step inherent to T1EPI normalization pipelines represents an added challenge, which, if not adequately addressed, can limit precision and accuracy. Although they nominally represent the same brain, the geometric distortions and signal loss typical of EPI data, and the differences in contrast between T2* and T1-weighted images render co-registration a complex, non-linear problem (Hutton et al., 2002; Jezzard, 2012). Simple rigid-body (6*-df*) co-registration techniques cannot fully address these artifacts, resulting in suboptimal co-registration accuracy. Ultimately, co-registration errors propagate to atlas-space, where they reduce precision and accuracy Several methods have been developed for correcting EPI geometric distortions and improving the accuracy of co-registration, including nonlinear registration of EPI data to a non-distorted image from another modality (e.g., Wang et al., 2017) as well as various distortion estimation (i.e., field mapping) and correction techniques (Andersson, Skare, & Ashburner, 2003; Chung et al., 2011; Cusack, Brett, & Osswald, 2003; Holland, Kuperman, & Dale, 2010; Hutton et al., 2002; Jezzard & Balaban, 1995). Although these methods can enhance the quality of coregistration (Cusack et al., 2003; Hong, To, Teh, Soh, & Chuang, 2015; Togo et al., 2017), they cannot recover data from regions of signal loss or compression and they cannot account for more complex interactions between distortion and subject motion (Andersson & Sotiropoulos, 2016; Barry et al., 2010; Graham et al., *in press*).

Calhoun and colleagues recently reported a systematic comparison of several EPIO and T1EPI spatial normalization pipelines, focusing on measures of precision only (Calhoun et al., 2017). Across several metrics and datasets, they provided evidence suggesting that their EPIO approach is consistently more precise than the non-linear, ‘unified segmentation’ T1EPI approach implemented in SPM12 (Ashburner & Friston, 2005) and the diffeomorphic T1EPI approach implemented in ANTS (Avants, Epstein, Grossman, & Gee, 2008; Avants et al., 2011; Avants et al., 2010). Calhoun and colleagues show that T1EPI approaches result in greater voxelwise variability of the spatially normalized images across participants, their primary precision metric. The largest difference in variability between EPIO and T1EPI methods manifested as a ring or ‘halo’ of high-variability voxels at the outer edge of the brain. Remarkably, reduced precision was found even for data processed using a combination of boundary-based registration (BBR; Greve & Fischl, 2009) with distortion correction and diffeomorphic normalization with ANTS, methods previously shown to exhibit superior performance. Capitalizing on standard techniques for estimating participant motion, Calhoun and colleagues computed a second precision index by concatenating normalized EPI volumes across participants and then estimating the amount of rigid-body alignment required to register the data to an arbitrary reference participant. Again, their preferred variant of the SPM EPIO pipeline proved more precise, as indexed by a lower average levels of residual alignment. Finally, Calhoun and colleagues showed that their EPIO approach was also associated with larger t-values in a widely used go/no-go paradigm, underscoring the potential ‘real-world’ statistical advantages of EPIO over T1EPI. Calhoun and colleagues conclude by arguing that EPIO *without* fieldmap-based distortion correction “provides results as good as or in some cases better than” (p. 5340) T1EPI *with* distortion correction, contrary to the working practices of most imaging groups. These claims are intriguing because they seem counterintuitive and because they indicate that a change in standard data acquisition and processing practices may be warranted.

Here, we re-visit these conclusions and extend them to other measures of precision and to measures of accuracy. For consistency with Calhoun et al., we examine the T1EPI and EPIO approaches implemented in the widely used SPM12 software package as well as the EPIO variant they recommend. We also examine a T1EPI pipeline developed by our group at the University of Maryland, which harnesses tools from several software packages, including AFNI, ANTS, FSL, and SPM12 (cf. Tillman et al., 2017). We begin by describing the dataset that we analyzed and each of the processing pipelines that we compare. Next, we address the issue of how best to quantify spatial normalization precision. We show that voxelwise variance, the primary measure of precision used by Calhoun and colleagues, is confounded by systematic differences in normalized brain volume between the processing pipelines and present some alternative metrics. In the following section, we describe several measures of spatial normalization accuracy, an important goal of spatial normalization that was not considered by Calhoun and colleagues. Finally, we assess the practical significance of the four different normalization approaches, focusing on a widely used emotional faces task. This task is particularly useful for evaluating precision and accuracy because it makes it possible to assess the influence of different spatial normalization pipelines on BOLD signal differences in both the amygdala—a subcortical region that is challenging to image given its proximity to areas of signal distortion and dropout (LaBar, Gitelman, Mesulam, & Parrish, 2001; Parrish, Gitelman, LaBar, & Mesulam, 2000)—and in face-sensitive cortical regions, such as the fusiform cortex, that are far removed from such artifacts. On the basis of these results, we make specific recommendations for assessing and applying spatial normalization in future fMRI research.

## METHOD

### Participants

A total of 61 adults between the ages of 21 and 35 years were recruited from the local community as part of a larger study and financially compensated for their time. All had normal or corrected-to-normal color vision, and self-reported the absence of lifetime alcohol or substance abuse-related problems, lifetime neurological symptoms, current psychiatric diagnosis or treatment, pervasive developmental disorder or very premature birth, or a medical condition that would contraindicate MRI. Twelve participants were excluded from analyses due to unusable T1-weighted or spin echo (SE) datasets (*n*=3), technical problems with the scanner (*n*=1), incidental neurological findings (*n*=2), inadequate behavioral performance (>2 SDs below mean hit rate; *n*=3), or excessive motion artifact (*n*=3; see below), yielding a final sample of 49 subjects (*M*=22.4 years, *SD*=2.5; 46.9% female). Participants provided informed written consent and all procedures were approved by the University of Maryland Institutional Review Board.

### Procedures

After obtaining consent, an overview of the experimental procedures was provided and the emotional-faces/places task was described. MRI-compatible audio ear buds (Sensimetrics S14; Sensimetrics Co. Gloucester, MA) fitted with noise reducing ear plugs (Hearing Components Inc. St Paul MN) were provided for communication and hearing protection. Before entering the scanner room, a handheld metal detector was used to verify the absence of potentially hazardous ferromagnetic objects. The participant was positioned supine in the scanner and foam inserts were used to minimize potential movement and to further attenuate scanner noise. Visual stimuli were digitally back-projected onto a screen mounted at the head-end of the scanner bore and viewed using a mirror mounted on the head-coil. Participant status was continuously monitored from the control room using an MRI-compatible eye-tracker (data were not recorded; Eyelink 1000; SR Research, Ottawa, Ontario, Canada).

### Emotional-Faces/PIaces Paradigm

T2*-weighted EPI data were acquired while subjects performed a simple, fMRI-optimized, continuous-performance, emotional faces task. Building on work by our group (Stout, Shackman, Pedersen, Miskovich, & Larson, 2017) and many others (Hariri et al., 2002; Paulus, Feinstein, Castillo, Simmons, & Stein, 2005; Swartz, Knodt, Radtke, & Hariri, 2015) demonstrating the utility of emotional face paradigms for probing amygdala reactivity, subjects viewed alternating blocks of either emotional faces (8 blocks) or places (9 blocks). Block length (~16.3 s) was chosen to maximize our power to detect a difference in the blood oxygen level-dependent (BOLD) signal elicited by the two conditions (Henson, 2007; Maus, van Breukelen, Goebel, & Berger, 2010). To maximize signal strength and homogeneity and minimize potential neural habituation (Henson, 2007; Maus et al., 2010; Plichta et al., 2014), each block consisted of 16 brief presentations of faces or places (~1.02 s/image) (for related approaches, see O’Craven & Kanwisher, 2000; Williams et al., 2015). During face blocks, subjects used an MRI-compatible, fiber-optic response pad (MRA, Washington, PA) to discriminate (i.e., two-alternative, forced choice) between threat-related (i.e., fearful; 75% trials) and emotionally neutral expressions (25% trials) presented in a pseudorandomized order (Costafreda, Brammer, David, & Fu, 2008; Lindquist, Wager, Kober, Bliss-Moreau, & Barrett, 2012; Stout et al., 2017; Whalen, 1998). This design choice was aimed at reducing monotony and minimizing potential habituation to the fearful expressions (Plichta et al., 2014). During place blocks, subjects discriminated between frontal facades of suburban residential buildings (i.e., houses; 75%) and urban commercial buildings (i.e., skyscrapers; 25%). Face stimuli were taken from prior work by Gamer and colleagues (Gamer, Schmitz, Tittgemeyer, & Schilbach, 2013; Gamer, Zurowski, & Biichel, 2010; Scheller, Büchel, & Gamer, 2012) and included standardized images of unfamiliar male and female adults displaying unambiguous fearful or neutral expressions. To maximize the number of models, images were derived from multiple well-established databases: Ekman and Friesen’s Pictures of Facial Affect (Ekman & Friesen, 1976), the FACES database (Ebner, Riediger, & Lindenberger, 2010), the Karolinska Directed Emotional Faces database (http://www.emotionlab.se/resources/kdef), and the NimStim Face Stimulus Set (https://www.macbrain.org/resources.htm). Colored images were converted to grayscale, brightness normalized, and masked to occlude non-facial features (e.g., ears, hair). Grayscale building stimuli were taken from prior work by Choi and colleagues (Choi, Padmala, & Pessoa, 2012, 2015).

### MRI Data Acquisition

MRI data were acquired using a Siemens Magnetom TIM Trio 3 Tesla scanner and 32-channel head-coil. Sagittal T1-weighted anatomical images were acquired using a magnetization-prepared, rapid-acquisition, gradient-echo (MPRAGE) sequence (TR = 1,900 ms; TE = 2.32 ms; inversion time = 900 ms; flip angle = 9°; number of slices = 192; sagittal slice thickness = 0.9 mm; in-plane = 0.449 × 0.449mm; matrix = 512 × 512; field of view = 230mm × 230mm). To enhance spatial and temporal resolution, a multi-band sequence was used to collect a total of 286 oblique-axial EPI volumes during a single scan of the emotional faces task (multiband acceleration = 6; TR = 1,000 ms; TE = 39.4 ms; flip angle = 36.4°; slice thickness = 2.2 mm, number of slices = 60; voxel size in-plane = 2.1875 × 2.1875 mm; matrix = 96 × 96). Images were collected in the oblique-axial plane (approximately −20° relative to the ACPC plane) to minimize susceptibility artifacts. To enable distortion correction, two oblique-axial SE images were collected in each of two opposing phase-encoding directions (anterior-to-posterior and posterior-to-anterior) at the same location and spatial resolution as the functional volumes (TR = 7,220 ms; TE = 73ms).

### MRI Data Processing

The major aim of this study was to compare the precision, accuracy and ‘real-world’ statistical performance of T1EPI and EPIO spatial normalization techniques. Building on work by Calhoun and colleagues, imaging data were processed using four pipelines:

- The standard T1EPI unified segmentation approach implemented in SPM12 (T1EPI_Standard_)
- The standard EPIO unified segmentation approach implemented in SPM12 (EPIO_Standard_)
- A variant of the SPM12 EPIO approach incorporating several recommendations from Calhoun and colleagues (EPIO_Calhoun_)
- The multi-tool T1EPI pipeline developed at the University of Maryland (T1EPI_Maryland_)

For assessment purposes, three kinds of output datasets were created for each pipeline:

- The temporal mean of the processed, spatially normalized, spatially **unsmoothed** EPI time-series for each participant
- The ‘mean-of-the-means’ or grand average of the temporal mean EPI images across participants
- The processed, spatially normalized, and spatially **smoothed** EPI time-series for each participant, which served as the input for whole-brain, voxelwise statistical analyses of the emotional-faces/places task

#### T1EPI_Standard_ Pipeline

For each participant, the first 3 volumes of each EPI dataset were removed to allow for signal equilibration. T1-weighted and functional EPI volumes were written into standard (RAI) orientation using *fslreorient2std* and the re-oriented EPI data were slice-time corrected to the TR midpoint using *3dTshift*. Remaining processing was performed using SPM12’s *preproc_fmri.m* script (ID 6177 2014-09-16) modified to avoid interactive file specification, utilize the MATLAB parallel toolbox, increase interpolation order, and alter the final bounding box and voxel dimension. Motion correction to the mean EPI volume in this script is implemented using the ‘Realign and Unwarp’ option. In keeping with Calhoun and colleagues and common practice, no distortion correction was applied at this step. Next, the script bias corrects and (simultaneously) segments and spatially normalizes the T1-weighted volume to the MNI atlas using the ‘unified segmentation’ option (Ashburner & Friston, 2005) and default tissue probability maps included with SPM12. The script then rigidly co-registers the mean motion-corrected EPI to the T1-weighted image. This co-registration transformation is then applied to the motion-corrected EPI data. Finally, the T1-to-template (‘unified segmentation’) transformation is applied to the co-registered, motion-corrected EPI data and the resulting transformed data is resliced using 5^th^-order splines to 2mm^3^ within the field-of-view defined by the MNI152 2-mm template (i.e., the ICBM 152 non-linear 6th-generation asymmetric average brain stereotaxic registration model, Grabner et al., 2006). The normalized images are smoothed (6-mm FWHM) for statistical analysis.

#### EPIO_Standard_ Pipeline

The standard SPM12 EPIO pipeline is broadly similar to T1EPI_standard_. Here, processing was performed using a version of SPM12’s *preproc_fmri_without_anat.m* script modified as above. In this script, instead of normalizing the T1-weighted image, the mean motion-corrected EPI volume is directly segmented and spatially normalized to the MNI template using ‘unified segmentation’ and this transformation is applied to the motion-corrected EPI data. Other details were identical to those detailed for the T1EPI_standard_ pipeline.

#### EPIO_calhoun_ Pipeline

EPIO_calhoun_ is similar to EPIO_Standard_. modified to adhere to the conclusions of Calhoun et al., (2017). Here, rather than using unified segmentation, normalization was performed with the ‘old normalization’ option (Ashburner, Andersson, & Friston, 1999) using a 45-mm cut-off, resulting in a 4 × 5 × 4 basis set, per the recommendations of Calhoun et al. Other details were identical to those detailed for the T1EPI_standard_ pipeline.

#### T1EPI_Maryland_ Pipeline

T1EPI_Maryland_ is conceptually similar to T1EPI_standard_, but employs tools from multiple software packages (Tillman et al., 2017). ***Overview***. As described in more detail below, each participant’s T1-weighted anatomical image was inhomogeneity-corrected, brain-extracted, and diffeomorphically normalized to a customized version of the MNI 1mm T1 template image. EPI data were de-spiked, slice-time corrected, distortion-corrected, co-registered, normalized via the T1, and spatially smoothed. ***Anatomical data***. T1-weighted images were written into standard (RAI) orientation and rescaled (0-1000) using *ImageMath*, inhomogeneity-corrected using N4 (Tustison et al., 2010), and brain-extracted using a consensus method (for similar approaches, see Meyer, Padmala, & Pessoa, 2017; Najafi, Kinnison, & Pessoa, 2017; Souza et al., 2017). In brief, prior work indicates that spatial normalization is enhanced by using a brain-extracted (i.e., ‘skull-stripped’) template and brain-extracted T1 images (Acosta-Cabronero, Williams, Pereira, Pengas, & Nestor, 2008; Fein et al., 2006; Fischmeister et al., 2013; Klein et al., 2010) but our experience has suggested that this is only the case for accurate brain extraction. To ensure consistently high-quality extractions, we implemented a multi-tool, consensus strategy. For each T1 image, four extraction masks were generated. Three masks were generated using *BSE* (Shattuck, Sandor-Leahy, Schaper, Rottenberg, & Leahy, 2001), *ROBEX* (Iglesias, Liu, Thompson, & Tu, 2011), and combining grey and white matter compartments obtained via SPM unified segmentation (Ashburner & Friston, 2005) with an updated tissue probability map (Lorio et al., 2016), respectively. The fourth mask was generated by spatially normalizing the whole T1-weighted volume (skull on) to the MNI152 T1 whole volume (skull on) template using a diffeomorphic approach (*SyN*) with a mutual information cost function (Avants et al., 2008; Avants et al., 2011; Avants et al., 2010; Iglesias et al., 2011). The inverse of this spatial transformation was then applied to a version of the 1-mm MNI152 brain mask distributed with FSL that was manually refined to remove extra-cerebral tissue. Next, a best-estimate extraction mask was determined by consensus, requiring agreement across three or more extraction masks. Using this consensus mask, each T1-weighted image was extracted and spatially normalized to the refined, brain-extracted 1-mm MNI152 template using *SyN*. In addition, brain-extracted T1 images were segmented using *FAST* (Zhang, Brady, & Smith, 2001) for use in EPI-to-T1 co-registration^3^, as described below. Each dataset was visually inspected before and after processing for quality assurance. Brain extraction failed for one subject. Here, the mask was semi-automatically created using a combination of the SPM and inverse ANTS methods detailed above, as well as masks from *BET2* (Jenkinson, Pechaud, & Smith, 2005; Smith, 2002) and *3dSkullStrip* (Cox, 1996). ***Fieldmap correction***. Co-planar SE images with reversed phase-encoding directions were written to standard orientation and combined to create undistorted SE images and fieldmaps using *topup* (Andersson et al., 2003; Smith et al., 2004). To mitigate extreme values, fieldmaps were converted to radians; de-spiked and median filtered (default parameters); thresholded (min=−375, max=475); and spatially smoothed (smooth2=5, smooth3=2) using a combination of *fugue* and *fslmaths*. A single image was created by averaging the undistorted SE images. This average image was then inhomogeneity-corrected using N4 and brain-extracted using *3dSkullStrip*. ***EPI data***. EPI datasets were written to standard orientation, de-spiked using *3ddespike*, and slice-time corrected to the middle of the TR using *3dtshift*. For co-registration of the functional and anatomical images, an average EPI image was created using two-pass motion correction in *3dvolreg*. The average image was simultaneously coregistered with the T1-weighted image and fieldmap-corrected using *EPI_reg.sh* with the boundary-based cost function (Greve & Fischl, 2009) and the previously created fieldmap, undistorted average SE image, inhomogeneity-corrected T1-weighted image, brain masked T1-weighted image, and T1-weighted image white matter (WM) compartment. Unpublished observations by our group indicate that even with smoothing and thresholding of the fieldmap, the resulting distortion-corrected EPI images often contain areas that appear ‘over-corrected;’ particularly in midline regions of the medial temporal lobe and orbitofrontal cortex. In order to explicitly control the smoothness of the distortion correction as well as to create an ITK-compatible version of the distortion-correction warp field, we used ANTs to spatially transform the above pr*e-EPI_reg.sh* average EPI image directly to the co-registered, distortion-corrected EPI image output from *EPI_reg.sh*. The time-series images were then motion-corrected by rigidly aligning each volume to the average image using *AntsMotionCorr* and the estimated affine transformation matrix was saved for each image. To assess residual motion artifact, the variance of volume-to-volume displacement of a selected voxel in the center of the brain (x=5, y=34, z=28) was calculated using the motion-corrected EPI data. Subjects (n=3) with extreme motion variance (>2SDs above the mean) were excluded from analyses. At this point, the complete series of spatial transformations for each volume is known and was applied in a single step to avoid introducing additional spatial blurring. Thus, for each slice-time corrected image, the following spatial transformations were concatenated and applied in the following order: motion-correction rigid-body, co-registration-unwarping affine, co-registration-unwarping nonlinear, brain-extracted T1 affine, and brain-extracted T1 diffeomorphic. The transformed EPI images were resliced to 2-mm^3^ (5th-order splines) and spatially smoothed (6-mm FWHM) within the brain mask using *3dBlurInMask*. Importantly, other than the specific set of tools used for brain extraction and the intensity threshold values used to smooth the fieldmaps, no other parameters for any step in the T1EPI_Maryland_ were optimized for this specific data set. All other parameters (e.g., for *AntsMotionCorr, SyN, N4, EPI_reg.sh* etc.) were selected based evaluations on other similar datasets from multiple scanners.

#### Additional Post-Normalization Processing

For each combination of pipeline and participant, the temporal mean of the spatially normalized, spatially unsmoothed EPI time-series for each participant was segmented into gray matter (GM) and white matter (WM) compartments using SPM12 ‘unified segmentation’ and the default tissue prior map. Surface identifying brain masks were also created for the average images using *3dSkullStrip*. For each pipeline, the grand average of the temporal mean EPI images was created using *fslmerge* and *fslmaths*.

### Precision Metrics

The primary performance metric used by Calhoun et al., voxelwise variability, is biased. Voxelwise variability assumes an equivalent normalization target; that is, it assumes that all of the images from all of the pipelines have been transformed into the identical stereotaxic space with no systematic error. In principle, this assumption is reasonable because all of the pipelines nominally target the MNI152 6^th^-generation atlas. The problem is that there is a consistent misalignment between images normalized via EPIO_calhoun_ and the other three pipelines; EPIO_calhoun_ normalized images are consistently smaller in the anterior-to-posterior (AP) and left-to-right (LR) dimensions compared to the other pipelines.

To demonstrate this visually, we compared the *3dSkullStrip* brain masks for each pipeline (see above) to one another and to the 2-mm MNU52 T1-weighted template. The masks provide a means of identifying voxels lying on or intersecting with the outer surface of the brain (i.e. mask values 2,3, and 4). We created surface-intersecting masks (‘surfaces’) for each participant and generated a sum of these surfaces across participants for select pipelines. **Figure 1** shows the resulting surfaces for the EPIO_calhoun_ and T1EPI_standard_ pipelines, with the 2-mm MNI152 T1-weighted template shown for comparison. The surface of the EPIO_calhoun_ brain consistently falls inside the surface of the MNI152 template. Subtracting the two shows that the EPIO_calhoun_ surface is substantially smaller with a ‘halo’ of T1EPI_standard_ surface around the EPIO_calhoun_ surface, mirroring the report of Calhoun and colleagues.

**Figure 1.**
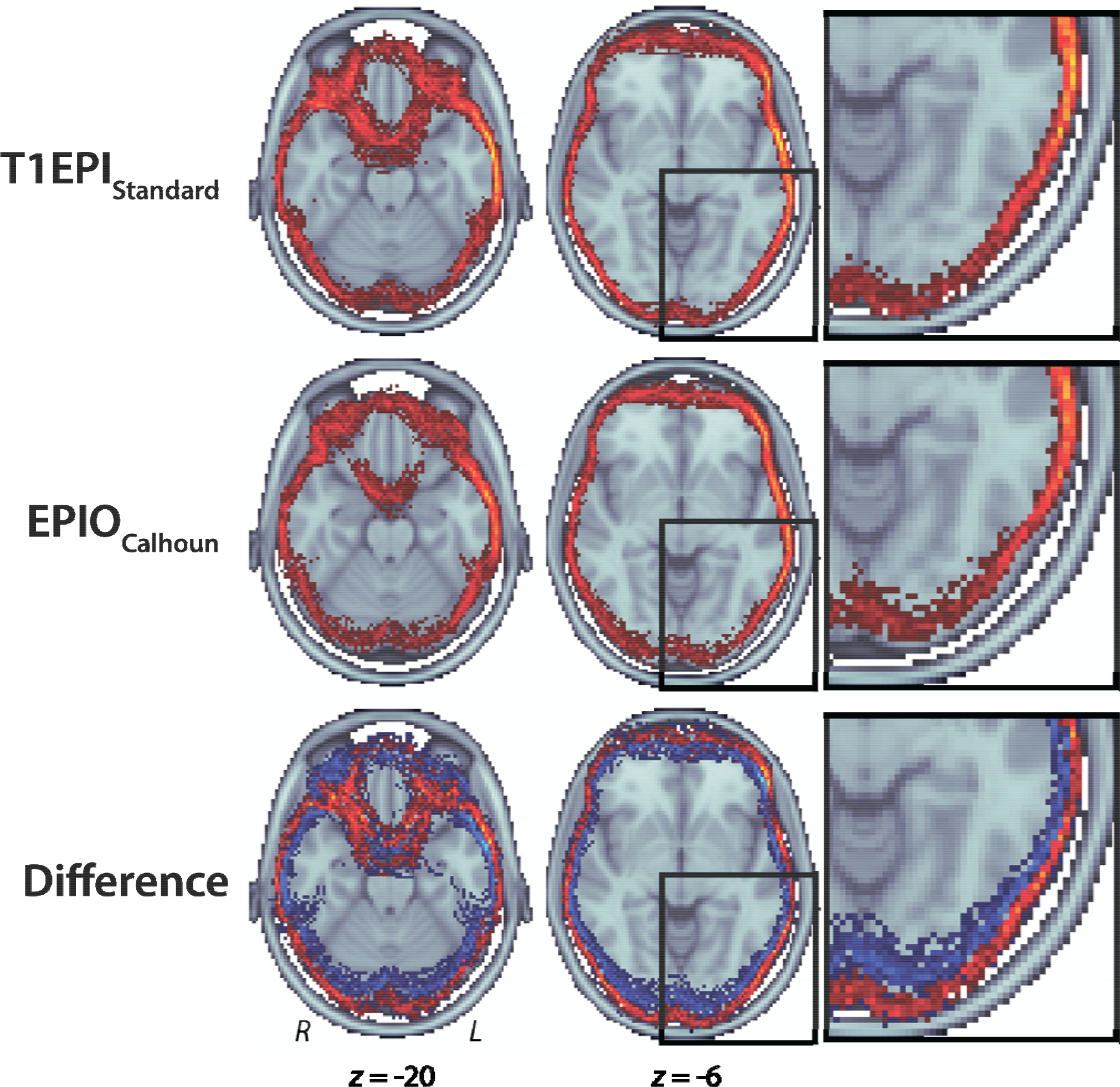
Brain ‘surfaces’ produced by the T1EPI_Standard_ and EPIO_calhoun_ pipelines superimposed on the 2-mm T1-weighted MNI template for representative axial slices. Inset panels depict a magnified view of the left-posterior surface. Surface-intersecting masks (‘surfaces’) were created, summed across participants, and mean thresholded for each pipeline (top and middle rows). Brighter colors indicate greater overlap. The surface of the EPIO_calhoun_ brain consistently falls inside the edge of the template brain. As shown in the bottom row, the difference image (T1EPI_Standard_ minus EPIO_calhoun_) shows that the EPIOCalhoun surface is substantially smaller [*blue*, negative values) with a ‘halo’ of T1EPI_Standard_ surface [*red*, positive values) at the edge of the template brain, mirroring the observations of Calhoun and col-leagues (their figures 3-5).

To quantitatively assess the difference, we estimated surface extent by subtracting the minimal AP and LR surface coordinate from the maximal for each participant. Consistent with **Figure 1**, the mean EPIO_calhoun_ surface was significantly smaller than the mean T1EPI_standard_ surface in both the AP and XY dimensions, *ts*(48) > 4.20, *ps* < .001 (EPIO_calhoun_, AP: *M* = 175.68 mm, *SE* = 3.50; XY: *M* = 140.82 mm, *SE* = 1.96; T1EPI_standard_, AP: *M* = 184.08 mm, *SE* = 4.40; XY: *M* = 142.28 mm, *SE* = 2.62). For perspective, we calculated the surface extent of the MNI152 2mm T1-weighted template using the same methods (MNI152: AP = 182 mm, XY = 143 mm). Thus, relative to the MNI152 T1 template the EPIO_calhoun_ surface extent was >6 mm smaller than the MNI surface extend in the AP dimension and >2 mm smaller in the LR dimension on average. In contrast, the T1EPI_standard_ surface extent was ~2 mm larger in the AP dimension and <1 mm larger in the LR dimension than the template. Similar results were obtained for analyses performed using extent measures derived using the GM and WM compartments or 25% maximum intensity surface (not reported). In short, voxelwise variability—one of the core metrics used by Calhoun and colleagues—cannot be used to compare the precision of different spatial normalization pipelines, at least near the brain’s surface, where it confounds intra-and extra-cerebral voxels. To circumvent this limitation, we developed six alternative measures of precision, each endowed with different strengths and weaknesses.

### Grand average spatial smoothness

For descriptive purposes, we estimated the spatial smoothness of the grand-average EPI images using *3dFWHM*. The underlying logic is similar to voxelwise variability, but sidesteps potential differences in cerebral extent. To the extent that all of the participants’ EPI images are perfectly aligned with one another, the grand-average image will have the same degree of smoothness as that of the individual images, whereas imprecise alignment will result in blurring and increase the smoothness of the grand average image. For this metric, decreased smoothness indicates superior precision.

### Grand average correlations

To the extent that the individual EPI data are in perfect alignment, voxelwise correlations between each participant’s mean EPI volume and the grand average will approach unity. We used MATLAB (R2017a; Mathworks Inc. Natick MA), to compute the voxelwise correlations. The resulting correlation coefficients were Fisher *R*-to-*Z* transformed for statistical comparison. While perhaps the simplest measure, when computed over the entire volume, such correlations may be misleading because they reflect a mixture of higher intensity cerebral and extra-cerebral voxels as well as the low intensity, empty background (i.e., a bimodal distribution of voxelwise signal values), likely violating the assumption of normally distributed residuals. To mitigate this potential concern as well as to avoid influence of signals in unimportant tissue, individual to grand average correlations were also calculated using a subset of cerebral voxels. To this end, an unbiased brain mask was created by twice eroding the 2-mm brain mask distributed with FSL. For both the whole-volume and intracerebral analyses, stronger correlations indicate superior precision.

### Inter-participant correlations

The grand average correlation metric described above can be influenced by the smoothness of the grand average itself. Differential information (e.g., increased smoothness) in the grand average image could impact the magnitudes of the correlations yielding estimates of precision that may not be directly comparable across pipelines. Therefore, we selected the participant closest to the centroid of each pipeline (i.e., the individual showing the strongest correlation with the grand average) and then computed voxelwise correlations between the remaining participants’ EPI datasets and the reference participant. The three SPM-based methods identified the same individual (participant 34) as the reference. The T1EPI_Maryland_ pipeline identified a different individual (participant 16) as the reference. Therefore, separate analyses were computed using each of the references. As with the grand average correlations, inter-participant correlations were transformed and the analyses were repeated using the subset of cerebral voxels. As before, stronger correlations indicate superior precision.

### Affine correction to grand average

Building on the rigid-body (6-*df*) approach used by Calhoun and colleagues, *AntsMortionCorr* (mutual information cost function) was used to compute an affine (12-*df*) registration between each participant’s mean EPI and the grand average. Residual variability was then summarized by deforming 1,000 random points on a 50-mm sphere using the estimated affine matrices and computing the mean Euclidean displacement for each participant. Because this metric measures precision in real physical units (millimeters of average residual displacement of points on a standardized surface), it provides a measure of precision that can be directly compared across studies. Lower affine displacement distance indicates superior precision.

### Inter-participant S0rensen-Dice coefficient

Correlation-based metrics can be misled by image intensity differences that are not related to the underlying anatomy (e.g., differential task responses, image inhomogeniety etc.) and thus not related to normalization precision. To mitigate this potential issue, we examined an alternative measure of precision based on the degree of binary spatial overlap of anatomy across individual EPI datasets. We compared the overlap of the WM compartments identified by SPM 12 ‘unified segmentation’ for each participant’s average EPI image using the Sørensen-Dice coefficient. We focused on the WM compartment because it proves a robust means of identifying the underlying brain anatomy for each individual. Potential alternative anatomical markers are more problematic. For example, hand-drawn landmarks are difficult to identify consistently in low resolution EPI and it is more challenging to consistently distinguish GM from the neighboring cerebrospinal fluid (CSF) compartment in T2*-weighed images. We thresholded (*p* > .80) and binarized the native space WM probability images and then used the Sørensen-Dice coefficient to quantify the degree of overlap between each participant’s WM compartment and that of the two reference participants (subjects 34 and 16; see the prior section for details). We eschewed comparison with the grand average images, due to the poor quality of attempted tissue segmentations on these images (not reported). Higher Sørensen-Dice coefficients indicate superior precision.

### Surface Agreement

A final, essentially nominal, normalization precision metric is the inter-participant overlap of the previously described brain surfaces (generated from the *3dSkullStrip* brain masks). We binarized the masks into surface and non-surface as before and then summed the voxels across participants. Given perfect normalization precision at the surface, all voxels would take values of zero or 49 (the number of participants). We report the mean of the non-zero voxels for each pipeline. Although a *t* test on these means could be performed, the sheer number of observations practically guarantees statistical significance, rendering the test uninformative (Killeen, 2005).

### Accuracy Metrics

An accurate spatial normalization pipeline is one that minimizes the amount of residual variability between the individual functional datasets and the target atlas, here, the 6th-generation MNI152. The accuracy of EPI-to-MNI152 normalization cannot be quantified using correlations because of marked differences in tissue contrast. In this section, we describe several alternative measures. In all cases, the 2mm T1-weighted MNI152 template, which was not directly used in any of the normalization pipelines, served as the reference.

### Mutual Information (MI) with template

We estimated the degree of MI between each participant’s mean EPI image and the template using a bivariate histogram approach implemented in MATLAB. As before, this was computed separately for both the whole volume as well as the subset of cerebral voxels. For the subset here, we simply used the standard 2-mm MNI152 brain mask distributed with FSL. Because this mask is already a fairly faithful representation of the target template, no additional erosion was necessary. Higher MI indicates superior accuracy.

### Participant-Template Sørensen-Dice coefficient

As above, the Sørensen-Dice coefficient was computed by comparing the WM compartment derived from each participant with that derived from the template. Higher Sørensen-Dice coefficients indicate superior accuracy.

### Participant-Template BBR cost

The use of SPM ‘unified segmentation’ to estimate the Participant-Template Sørensen-Dice coefficient introduces a potential bias, one that most favors the EPI_standard_ pipeline. To circumvent this, we used *fast* (Zhang et al., 2001) to segment the 2-mm T1-weighted MNI152 template distributed with FSL and then computed the BBR cost between each participant’s mean EPI image and the template. The BBR cost function identifies the boundaries of the WM compartment using the T1-weighted template and then computes the intensity gradient of the EPI along a normal to the boundary (Greve & Fischl, 2009). Even with the limited contrast afforded by T2*-weighted images, the image intensity should vary in the correct direction across the boundary line. It merits comment that, while BBR is part of the T1EPI_Maryland_ pipeline that aligns the participant’s EPI to their T1-weighted image, it is not used to compute the spatial normalization, that is, to compute the T1-weighted image to template transform. BBR with the template is never used in the T1EPI_Maryland_ pipeline. However, to the extent that BBR reduces co-registration error that would otherwise propagate to atlas space, this metric is biased to favor the T1EPI_Maryland_ pipeline. Lower BBR costs indicate superior accuracy.

### ‘Real-World’ Performance: Statistical Parametric Mapping

Building on the work of Calhoun and colleagues, we also assessed the practical or ‘real-world’ statistical significance of using a particular spatial normalization pipeline. Here, SPM12 was used to quantify BOLD signal changes in response to the emotional-faces/places paradigm. At the first level, the task was modeled using a simple boxcar function convolved with a canonical hemodynamic response function, with place blocks serving as the implicit baseline. Temporal autocorrelation was modeled using the ‘FAST’ option. Nuisance variates included 6 motion-related estimates. The main effect of Condition was assessed using the resulting first-level contrast images and a voxelwise one-sample *t* test. Results are reported for voxels surviving a *p*<.05, whole-brain corrected for familywise error (FWE), threshold. We primarily focus on results in the vicinity of bilateral amygdala and fusiform gyrus due to their involvement in the emotional faces task as well as their presumed differential susceptibility to EPI distortion. Following Calhoun and colleagues, we report the differences in ‘effective number of participants’ (*N*_*eff*_) for each pipeline, where *N*_*eff*_ represents the number of additional participants required to achieve a comparable *t* value.

### Post-hoc Analysis: Relation between precision and group level t statistics

Using the magnitude of second-level statistical parameters (i.e., *t* or *N*_*eff*_) as a ‘real-world’ evaluation of precision assumes that these values scale with the precision of spatial normalization (cf. Calhoun et al., 2017) and are not be driven by other factors. To better evaluate this, we examined the distance between activation peaks identified in single-subject data and those identified in the overall second-level group results (see below). For each combination of participant and pipeline, the SPM *t* image was thresholded at *p*<0.10, uncorrected. All local maxima in this thresholded image were identified using *spm_get_lm*. Local maxima were sorted in descending order by *t* and those within 8 mm of a maximum with a larger *t* were discarded as redundant. For each peak identified in the overall group results, the Euclidian distances to the remaining local maxima were evaluated and the distance of the closest maximum recoded. The mean of these distances across participant was evaluated for each combination of participant, pipeline, and peak. The resulting pipeline averages were centered (mean subtracted) within each peak. Likewise, the *t* values from the seven local maxima identified in the second-level SPMs for each pipeline were centered within peak and the resulting values compared to the centered mean distances. We also compared the distances across participants between the pipelines.

## RESULTS

### Precision

Summary statistics for the six measures of normalization precision are provided in **Table 1**. See the Supplementary Results for complete pairwise statistical comparisons.

**Table 1.** Precision: Pipeline Nominal Ranks and Metric Values.

| Precision Metric |  | Pipeline Nominal Rank Order and Value |  |  |  |
| --- | --- | --- | --- | --- | --- |
|  |  | T1EPI<sub>Standard</sub> | EPIO<sub>Standard</sub> | EPIO<sub>Calhoun</sub> | T1EPI<sub>Maryland</sub> |
|  | Grand Average Spatial Smoothness, mm (ranked from most to least smooth) | Third<br>39.792 | Fourth<br>40.281 | Second<br>39.307 | First<br>30.29 |
| Whole Volume | Grand Average Correlations | Fourth<br>0.937 | Third<br>0.939 | Second<br>0.943 | First<br>0.947 |
|  | Inter-Participant Correlations, Reference Subject 34 | Fourth<br>0.898 | Second<br>0.901 | Third<br>0.901 | First<br>0.905 |
|  | Inter-Participant Correlations, Reference Subject 16 | Fourth<br>0.891 | Third<br>0.898 | Second<br>0.9 | First<br>0.91 |
| Cerebral Voxels | Grand Average Correlations | Second<br>0.853 | Third<br>0.839 | Fourth<br>0.832 | First<br>0.862 |
|  | Inter-Participant Correlations, Reference Subject 34 | Second<br>0.739 | Third<br>0.722 | Fourth<br>0.701 | First<br>0.74 |
|  | Inter-Participant Correlations, Reference Subject 16 | Second<br>0.767 | Third<br>0.75 | Fourth<br>0.737 | First<br>0.78 |
|  | Affine Correction to Grand Average | Second<br>1.138 | Third<br>1.205 | Fourth<br>1.365 | First<br>1.062 |
|  | Inter-participant Sørensen-Dice coefficient, Reference Subject 34 | Second<br>0.78 | Third<br>0.776 | Fourth<br>0.751 | First<br>0.846 |
|  | Inter-participant Sørensen-Dice coefficient, Reference Subject 16 | Second<br>0.787 | Third<br>0.781 | Fourth<br>0.763 | First<br>0.853 |
|  | Surface Agreement, *n* | Fourth<br>14.26 | Third<br>14.67 | Second<br>14.85 | First<br>15.67 |

#### Grand average spatial smoothness

As seen in **Figure 2**, the T1EPI_Maryland_ pipeline produced the sharpest (i.e., least smooth) grand average image, followed by EPIO_calhoun_, T1EPI_standard_, and finally EPIO_Standard_, although the spatial smoothness of the latter three is quite similar (**Table 1** and **Supplementary Table 1**). The grand average EPI image produced using the T1EPI_Maryland_ pipeline is 1.87mm (FWHM) sharper than that created using the EPIO_calhoun_ pipeline, which in turn, is only 0.25-mm sharper than those produced by the T1EPI_standard_ and EPIO_Standard_ pipelines. Converting the global FWHM to standard deviations, the smoothness of the T1EPI_Maryland_ grand average is ~3.74-mm, while the remaining images exceed 4.53-mm.

**Figure 2.**
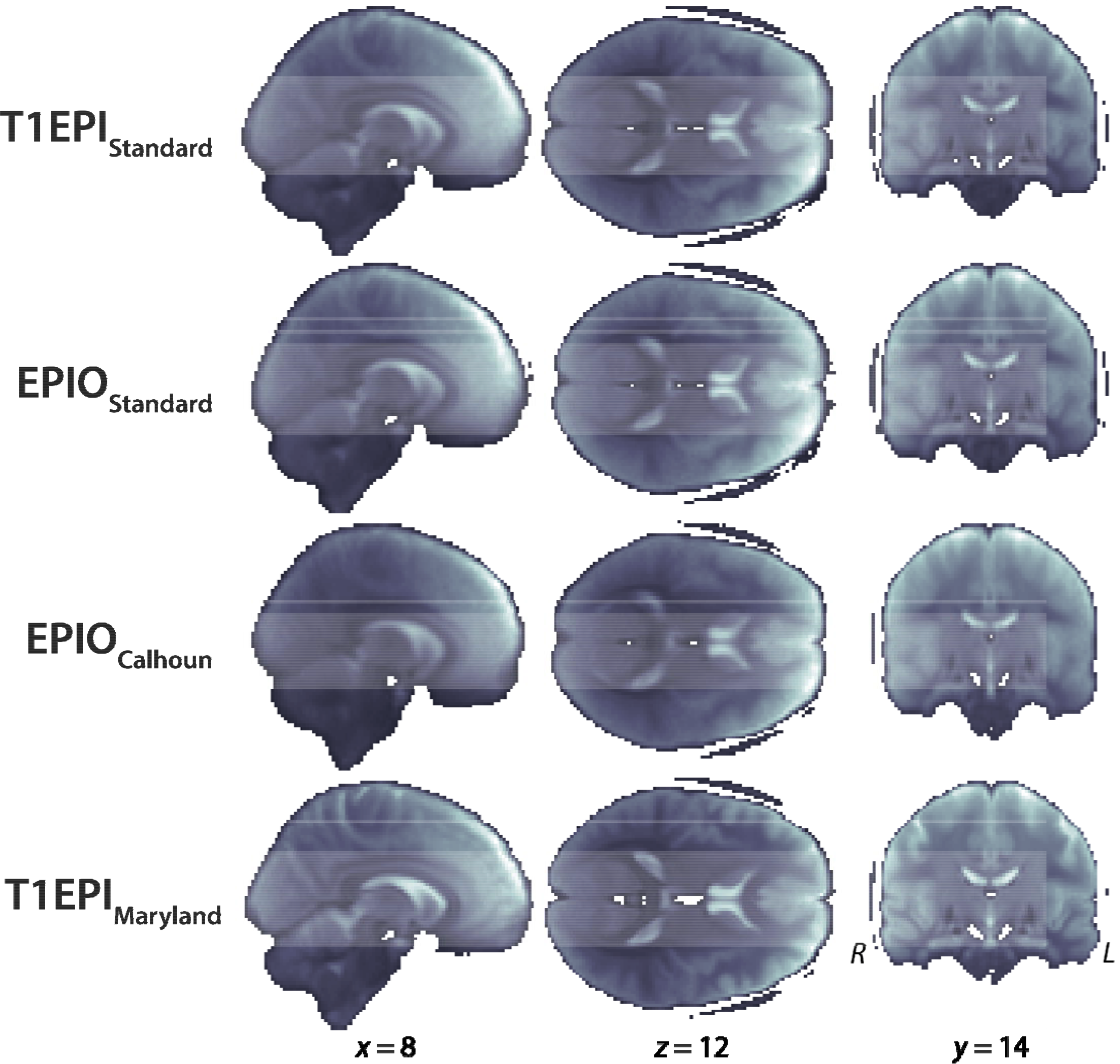
Grand average EPI images produced by each spatial normalization pipeline. Visual inspection suggests systematic differences in the sharpness of many major anatomical landmarks (e.g., cingulate sulcus, Heschl’s gyrus, lateral ventricles, parieto-occipital fissure). Overall differences in cerebral shape are also apparent.

#### Grand average correlations

Considering the whole volume, the T1EPI_Maryland_ pipeline produced the strongest correlations between the single-subject and grand average images, followed by EPIO_calhoun_, EPIO_Standard_, and finally T1EPI_standard_. All of the pairwise comparisons between pipelines were significant (*ts*(48) > 2.35, *ps* < .03) except for the difference between EPIO_Standard_ and T1EPI_standard_ pipelines (*p* = .07) (see **Supplementary Table 2**). Within the subset of cerebral voxels, the results were somewhat different. Again, images from the T1EPI_Maryland_ pipeline were most strongly correlated with the corresponding grand average, now followed by T1EPI_standard_, EPIO_Standard_, and finally EPIO_calhoun_. All of the pairwise comparisons were significant, *ts*(48 > 3.44, *ps* < .002 (see **Supplementary Table 3**).

#### Inter-participant correlations

Considering the whole volume, the T1EPI_Maryland_ pipeline produced the strongest correlations between the individual participant’s images and the single-subject reference image. When participant 34 served as the reference, T1EPI_Maryland_ was followed by EPIO_Standard_, EPIO_calhoun_, and finally TIEPIO_Standard_. When participant 16 was the reference, T1EPI_Maryland_ was followed by EPIO_calhoun_, EPIO_Standard_, and finally TIEPIO_Standard_. The significance of particular pairwise comparisons differed somewhat depending upon reference participant (see **Supplementary Tables 4-5**)^4^. Within the eroded brain mask, the T1EPI_Maryland_ pipeline again showed the strongest performance, followed by T1EPI_standard_, EPIO_Standard_, and finally EPIO_calhoun_. This pattern was independent of reference participant, although the significance of some pairwise comparisons again differed slightly (**Supplementary Tables 6-7**)^5^.

#### Affine correction to grand average

Images from the T1EPI_Maryland_ pipeline required the least affine (12-*df*) correction to match the corresponding grand average, followed by those from the T1EPI_standard_, EPIO_Standard_, and finally the EPIO_calhoun_ pipeline. The T1EPI_Maryland_ and T1EPI_standard_ pipelines both required significantly less correction than the EPIO_calhoun_ pipeline, *ts*(48) > 2.38, *ps* < 0.03 (**Supplementary Table 8**). None of the other pairwise differences were significant.

#### Inter-participant Sørensen-Dice coefficient

The T1EPI_Maryland_ pipeline showed the most and the EPIO_calhoun_ pipeline showed the least overlap—as indexed by the Sørensen-Dice coefficient for the binarized WM compartments—and this was consistent across the choice of reference participant. For both references, images from the T1EPI_Maryland_ pipeline consistently outperformed those from the other three pipelines (*ts*(47) > 7.29, *ps* < .001) (**Supplementary Tables 9-10**). Likewise, images from the T1EPI_standard_ and EPIO_Standard_ pipelines consistently outperformed those produced using the EPIO_calhoun_ pipeline, *ts*(47) > 4.85, *ps* < .001. The relative ranking of the T1EPI_standard_ and EPIO_Standard_ pipelines, however, was conditional on the choice of reference participant. Using participant 34 as the reference, T1EPI_standard_ outperformed EPIO_Standard_ (*t*(47) = 13.57, *p* < .001), whereas the reverse pattern was observed for participant 16 (*t*(47) = 3.68, *p* < .001).

#### Surface Agreement

The T1EPI_Maryland_ pipeline showed the best inter-participant surface agreement (*M* = 15.67, *SE* = 0.045) followed by EPIO_calhoun_ (*M* = 14.85, *SE* = 0.041), EPIO_Standard_ (*M* = 14.67, *SE* = 0.042), and finally T1EPI_Standard_ (*M* = 14.26, *SE* = 0.033).

### Accuracy

Summary statistics for the three measures of accuracy are provided in **Table 2**. See the **Supplementary Results** for complete pairwise statistical comparisons.

**Table 2.** Accuracy: Pipeline nominal ranks and metric values.

| Accuracy Metric | Pipeline Nominal Rank Order and Value |  |  |  |
| --- | --- | --- | --- | --- |
|  | T1EPI<sub>Standard</sub> | EPIO<sub>Standard</sub> | EPIO<sub>Calhoun</sub> | T1EPI<sub>Maryland</sub> |
| Mutual Information (MI)<br>with Template,<br>Whole Volume | Third<br>**0.441** | Second<br>**0.445** | Fourth<br>**0.433** | First<br>**0.483** |
| Mutual Information (MI)]<br>with Template,<br>Cerebral Voxels | Third<br>**0.096** | Second<br>**0.099** | Fourth<br>**0.085** | First<br>**0.134** |
| Participant-Template<br>Sørensen-Dice Coefficient | Second<br>**0.714** | Third<br>**0.694** | Fourth<br>**0.669** | First<br>**0.725** |
| Participant-Template<br>BBR cost | Second<br>**0.639** | Third<br>**0.719** | Fourth<br>**0.766** | First<br>**0.510** |

#### MI with template

Images from the T1EPI_Maryland_ pipeline were most similar to the 2-mm T1-weighted MNI152 template, followed by those from the EPIO_Standard_, T1EPI_Standard_, and finally the EPIO_calhoun_ pipeline. This pattern was consistently observed, whether we focused on the whole volume or the subset of cerebral voxels. All pairwise comparison were significant, *ts*(48) > 2.57, *ps* < .02 (**Supplementary Tables 11-12**).

#### Participant-Template S0rensen-Dice coefficient

On average, the T1EPI_Maryland_ pipeline showed the most overlap with the MNI152 template WM compartment, followed by T1EPI_Standard_, EPIO_Standard_, and finally EPIO_calhoun_. All pairwise comparisons were significant, *ts*(48) > 6.87, *ps* < .001 (**Supplementary Table 13**).

#### Participant-Template BBR cost

On average, the T1EPI_Maryland_ pipeline was most accurate, as indexed by residual BBR cost, followed by the T1EPI_Standard_, EPIO_Standard_, and finally the EPIO_calhoun_ pipelines. All pairwise comparisons were significant, *ts*(48) > 13.39, *ps* < .001 (**Supplementary Table 14**).

#### ‘Real-World’ Performance: Statistical Parametric Mapping

Second-level statistical parametric maps are depicted in **Figure 3** and cluster statistics are provided in **Table 3**. Consistent with prior work, voxelwise regression analyses consistently identified significant BOLD signal changes (*p* < .05 whole-brain FWE corrected) in the bilateral amygdala and fusiform gyrus. Two other large clusters were consistently identified, one encompassing three maxima in the right visual cortex (posterior lateral occipital cortex [LOC] and superior temporal sulcus [STS]) and a second encompassing two maxima in the right prefrontal cortex (inferior and middle frontal gyri [IFG/MFG]). All locations were confirmed using the Harvard-Oxford cortical 2mm Atlas (Desikan et al., 2006)^6^.

**Figure 3.**
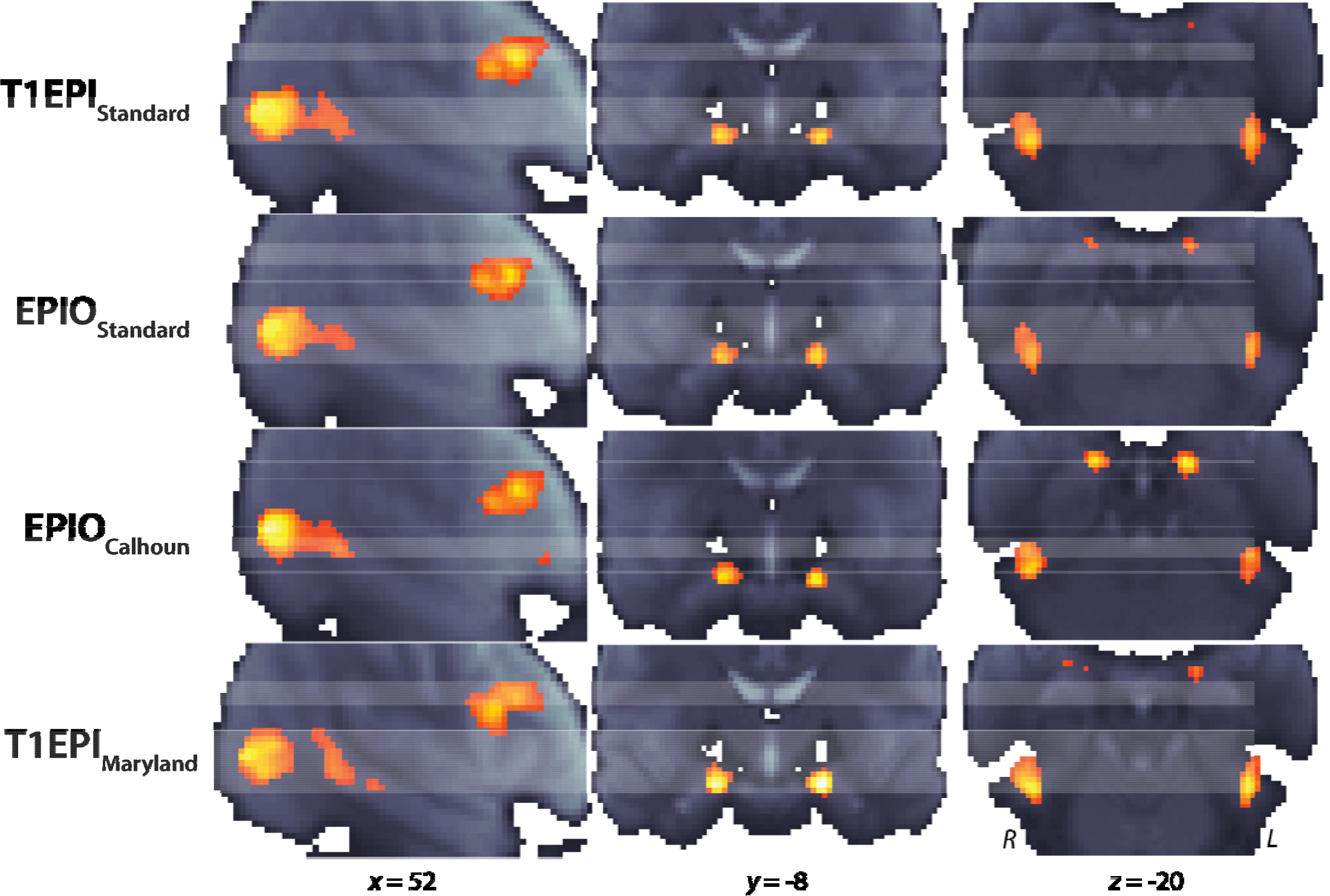
Second-level statistical parametric maps for the emotional-faces/places paradigm. Images depict the face-sensitive clusters (*p* < 0.05, whole-brain FWE corrected). Brighter colors indicate more positive *t* values. From left to right, significant clusters were observed in the right visual cortex, right lateral prefrontal cortex, bilateral amygdala, and bilateral fusiform cortex.

**Table 3.** Cluster and Maxima Statistics for the Emotional Faces/Places Paradigm.

|  | T1EPI<sub>Standard</sub> |  |  | EPIO<sub>Standard</sub> |  |  |
| --- | --- | --- | --- | --- | --- | --- |
| | Resels = 75.37 | Threshold ($p_{fwe} < 0.05$) = 5.83 | | Resels = 73.81 | Threshold ($p_{fwe} < 0.05$) = 5.83 | |
| | Peak MNI<sub>xyz</sub>(mm) | $t$ | Extent (voxels) | Peak MNI<sub>xyz</sub>(mm) | $t$ | Extent (voxels) |
| Left Amygdala | -18, -6, -16 | 8.97 | 55 | -18, -8, -16 | 8.69 | 44 |
| Right Amygdala | 20, -6, -14 | 9.38 | 68 | 20, -8, -18 | 8.55 | 57 |
| Left Fusiform | -42, -46, -20 | 8.08 | 106 | -42, -46, -22 | 8.33 | 94 |
| Right Fusiform | 44, -48, -20 | 8.67 | 138 | 44, -46, -24 | 8.84 | 130 |
| Right Visual Cortex | 54, -64, 8 | 10.98 | 619 | 54, -60, 8 | 10.45 | 526 |
|  | 50, -38, 0 | 6.73 |  | 46, -32, -2 | 6.59 |  |
|  | 56, -44, 4 | 6.91 |  | 50, -36, 2 | 6.47 |  |
| Right PFC | 50, 12, 20 | 8.1 | 392 | 50, 10, 20 | 8.32 | 336 |
|  | 52, 20, 24 | 9.18 |  | 52, 18, 24 | 8.88 |  |
|  | EPIO<sub>Calhoun</sub> |  |  | T1EPI<sub>Maryland</sub> |  |  |
| | Resels = 74.86 | Threshold ($p_{fwe} < 0.05$) = 5.83 | | Resels = 68.42 | Threshold ($p_{fwe} < 0.05$) = 5.78 | |
| | Peak MNI<sub>xyz</sub>(mm) | $t$ | Extent (voxels) | Peak MNI<sub>xyz</sub>(mm) | $t$ | Extent (voxels) |
| Left Amygdala | -16, -8, -20 | 9.85 | 46 | -20, -8, -16 | 12.5 | 83 |
| Right Amygdala | 18, -6, -20 | 8.89 | 61 | 22, -8, -16 | 11.49 | 90 |
| Left Fusiform | -42, -42, -22 | 7.82 | 94 | -40, -50, -18 | 10.11 | 132 |
| Right Fusiform | 44, -48, -20 | 8.51 | 117 | 42, -48, -20 | 9.58 | 178 |
| **Right Visual Cortex** | 54, -62, 10 | 11.51 | 548 | 54, -68, 6 | 10.72 | 593 |
|  | 52, -38, 4 | 7.13 |  | 54, -40, 2 | 7.22 |  |
|  | 56, -42, 10 | 6.78 |  | 58, -52, 8 | 7.17 |  |
| **Right PFC** | 50, 10, 20 | 8.2 | 342 | 54, 14, 22 | 9.93 | 403 |
|  | 52, 20, 24 | 8.92 |  | 54, 20, 26 | 8.3 |  |

Within the two *a priori* regions-of-interest (ROIs), the bilateral amygdala and fusiform cortex, the T1EPI_Maryland_ pipeline produced consistently stronger effects than the other pipelines, equivalent to a pairwise increase in *N*_*eff*_ of 17.1-104.7% (*M* = 56.9%, *SE* = 27.2). Among the three remaining SPM-based pipelines only, differences were conditional on ROI (mean nominal ranks across the four ROIs: T1EPI_standard_: 2.75, EPIO_Standard_: 3.00, EPIO_calhoun_: 3.25; **Table 3**). T1EPI_standard_ produced stronger effects in the right amygdala (*N*_*eff*_ increase of 14.0-19.9%); EPIO_calhoun_ produced stronger effects in the left amygdala [*N*_*eff*_ increase of 20.2-27.9%); and EPIO_Standard_ pipeline produced stronger effects in the fusiform ROIs (*N*_*eff*_ increase of 7.8-15.7%).

Outside of the *a priori* ROIs, performance was even more variable (mean nominal ranks across the five remaining peaks: T1EPI_standard_: 2.40, EPIO_Standard_: 3.40, EPIO_calhoun_: 2.20, T1EPI_Maryland_: 2.00). For example, within the right visual cortex cluster, data processed using T1EPI_Maryland_ produced the strongest effects in the two rostral maxima (*x* = 51, *y* = −37, *z* = 1 and *x* = 55, *y* = −44, *z* = 6; *N*_*eff*_ increases of 2.5-22.3%), whereas that processed using EPIO_calhoun_ produced the strongest effects in the caudal maximum (*x* = 54, *y* = −64, *z* = 8; *N*_*eff*_ increase of 9.7-17.1%)^7^.

#### Post-hoc Analysis: Relation between precision and group level t statistics

Uncentered mean distances between participant local maxima and group level peaks are shown in **Table 4**. Data processed with the T1EPI_Maryland_ pipeline was nominally less variable in local maximum distances compared to the other pipelines in every region except the visual cortex (i.e., STS). When centered within region, the degree of spatial dispersion was strongly and negatively related to the magnitude of regionally centered *t* statistics in the second-level SPMs (*r* = −0.654, *p* < 0.001), indicating that reduced spatial dispersion (i.e., increased precision) generally improves real-world results. When we compared pipelines, data processed using the T1EPI_Maryland_ pipeline showed significantly less dispersion on average compared to that processed using the three SPM-based pipelines (*ts* > 2.73, *ps* < 0.034; centered distance data). No other pairwise between pipelines were significant (*ts* < 0.82, *ps* > 0.445).

**Table 4.** Dispersion of Single-Subject Local Maxima around Second-Level (Group) Peaks (mm).

| Local Maxima to Group Peak Distances | T1EPI<sub>Standard</sub> | EPIO<sub>Standard</sub> | EPIO<sub>Calhoun</sub> | T1EPI<sub>Maryland</sub> |
| --- | --- | --- | --- | --- |
| **Left Amygdala** | 4.754 | 4.945 | 4.736 | 4.238 |
| **Right Amygdala** | 6.346* | 5.774* | 5.787* | 4.604 |
| **Left Fusiform** | 6.297* | 5.879 | 6.038 | 5.633 |
| **Right Fusiform** | 6.856* | 6.207 | 7.224* | 5.712 |
| **Right Visual Cortex 1** | 6.629 | 7.594 | 6.801 | 7.126 |
| **Right PFC 1** | 7.371* | 7.345* | 7.105 | 6.601 |
| **Right PFC 2** | 6.804 | 6.675 | 6.704 | 6.204 |

\* $p < 0.05$ difference with T1EPI<sub>Maryland</sub>

## DISCUSSION

Spatial normalization is a fundamental component of the vast majority functional neuroimaging studies with important consequences for data quality, interpretation, and reproducibility. Although a number of approaches have been developed, contemporary investigators typically rely on multi-step, nonlinear approaches that capitalize on T1-weighted ‘anatomical’ images. Here, we re-visited the performance of a variety of spatial normalization approaches using a comprehensive suite of metrics. Although none of these metrics is a perfect index of normalization quality, our results make it clear that voxel-wise variability, the primary metric used by Calhoun and colleagues—is confounded by between-pipeline differences in normalization accuracy. In the remainder of this section, we discuss our results in more detail and make specific recommendations for future research.

Across all metrics, the multi-tool T1EPI pipeline developed at the University of Maryland (T1EPI_Maryland_) was consistently ranked first in terms of both normalization precision and accuracy (**Tables 1** - **2**). The enhanced precision and accuracy of the Maryland pipeline generally resulted in superior ‘real-world’ statistical performance. This was found for both the magnitude of statistical effects at the group level in most regions (**Table 3**) as well as reduced dispersion of local maxima at the level of individual participants (**Table 4**). Collectively, these results provide clear evidence that investing in optimal normalization and other aspects of data preprocessing are worthwhile and can yield substantial benefits.

Consistent with Calhoun and colleagues’ observations, we found that when the whole volume—including extracerebral voxels—was considered, data processed using EPIO pipelines were more strongly correlated with both the grand average EPI and the reference participants than data processed using the T1EPI_standard_ pipeline. In addition, EPIO processed data showed better nominal precision (i.e., cerebral surface alignment) compared to data processed using the T1EPI_standard_ pipeline. Surface gray matter is important to many functional neuroimaging studies and the improvement over T1EPI_standard_ is intriguing. Our surface agreement metric can only identify overlap in the brain boundary and does not indicate whether or not the grey matter on that boundary represents the same anatomical location across participants. Understanding how different pipelines perform at the cortical surface is an important challenge for future research, perhaps incorporating additional measures of surface complexity (Zhou et al., 1999; Zilles et al., 1988, Thompson et al., 2005). Still, it is clear that enhanced whole-volume and surface precision come at the cost of reduced precision within the brain—EPIO_calhoun_ showed consistently lower precision, as indexed by correlations with the eroded mask, Sørensen-Dice coefficients, and residual affine motion^8^. This pattern indicates that the superior surface agreement of EPIO_calhoun_ relative to T1EPI_standard_ does not reflect superior agreement of anatomic regions on this surface.

In addition to reduced intra-cerebral precision, the data processed by the EPIO_calhoun_ method was consistently the least accurate. The reduced accuracy of this approach limits the validity of reported coordinates and curtails the use of several important interpretive tools, including pre-defined anatomical labels, probabilistic anatomical regions, and meta-analytic information. Therefore, for studies where such tools are important we cannot recommend the EPIO_calhoun_ pipeline.

The reduced accuracy of the EPIO_calhoun_ data likely reflects the inadequacy of the target template rather than the underlying logic of the method. The SPM EPI template was created by averaging images obtained from a mere 13 participants. The EPI images were transformed to the MNI152 atlas by warping each individual’s GM segment to an average GM image generated from the atlas. The average normalized image was then smoothed with an 8-mm (FWHM) kernel to produce the SPM EPI template. As a consequence, it lacks the anatomical detail of even the lowest-quality grand average images from the present study (**Figure 1**). The development of a higher resolution, non-linear EPI template would, in all likelihood, substantially improve the accuracy of the EPIO pipelines. Whether such a template can overcome marked individual and scanner-specific differences in signal loss and distortion is unclear. It may well be that study-specific EPI templates represent a more appropriate approach for EPIO pipelines. Indeed, studies have demonstrated superior EPIO performance using study-specific EPI templates (Huang et al., 2010).

The impact of spatial normalization for statistical analyses is perhaps the performance metric of greatest interest to practitioners. As a consequence of time and cost, fMRI studies are often underpowered, contributing to false negatives and reduced reproducibility (Fox, Lapate, Davidson, & Shackman, 2018; Munafò et al., 2017; Poldrack et al., 2017; Shackman & Fox, in press). If modest changes processing could substantially increase statistical power it would be of obvious benefit to all stakeholders, from investigators and funding agencies to clinicians and patients. Considering recent controversy regarding the validity and stability of fMRI methods (Bennett & Miller, 2010; Eklund, Nichols, & Knutsson, 2016), it is encouraging to see here that similar maxima of similar intensity were identified across the four pipelines (**Table 3** and **Figure 3**). Differences in location were within generally within 2-4 mm (which is consistent with the limited average affine variability) and the magnitudes of the observed *t* statistics within the *a priori* ROIs were all substantial. In the absence of ground truth, differences of the magnitude seen here do not license strong conclusions and we do not wish to overstate the significance of the observed results. All of the pipelines obtained a level of precision that would likely be considered adequate for many studies, particularly those employing lower resolution EPI data and more aggressive spatial smoothing, and all produced comparable, significant results.

Using T1EPI_Maryland_ data, peak statistical effects were consistently increased in the smaller clusters (i.e., amygdala and fusiform) relative to the other three pipelines. Logically, it is precisely these focused clusters where increased precision should have greatest impact and we expect this to be the major effect of precision on the group analysis. In the larger clusters with multiple foci (i.e., right visual and prefrontal cortices), using T1EPI_Maryland_ data either elevated sub-peaks at the cost of magnitude of the main focus (visual cortex) or swapped the rank ordering of peaks within the cluster (prefrontal cortex). Again, without clear ground truth, the effects in these larger clusters are challenging to unambiguously interpret.

Among the three SPM-based pipelines, our group statistical results do not warrant definitive recommendations; differences between these pipelines tended to be modest and inconsistent. Consider the four *a priori* ROIs, where each of the SPM-based pipelines occupied each of the rank order positions (2^nd^ through 4^th^) at least once. A pipeline that resulted in a relative increase in *N*_*eff*_ increase of 24.5% at one location resulted in a relative *N*_*eff*_ decrease of 13.1% relative to the same competitor at a different location. These results underscore the importance of considering a range of regions when drawing conclusions about the relative statistical performance of different imaging pipelines.

Finally, we note a fundamental limitation to further research on this question. While careful improvement to spatial normalization processing pipelines can yield significant benefit over generic approaches, there can be no single asymptotic performance for all cases because there, ultimately, there is no perfect normalization, no truth to maximally approximate. The one-to-one correspondence between voxels or points on a surface is only valid at a gross level (Stelzer et al., 2014; Turner, 2016; Turner & Geyer, 2014). Whole brain normalization methods such as those explored here will continue to be useful for many types of analyses in the forseeable future. However, we anticipate that normalization fucused on particular ROIs or structures (e.g., hippocampus: Yassa & Stark, 2009; Yassa et al., 2010) will become more prevelant particularly as the limits of spatial resolution of fMRI data are increased (Harel, 2012; Vu et al., 2017).

## Conclusions

Spatial normalization to a standard atlas space is an fundamental component of most functional neuroimaging studies, with implications for the alignment of participants, the reporting of meaningful stereotactic coordinates (which, of course, propagate to meta-analyses), the use of atlas-based anatomical priors, as well as statistical effect sizes and reproducibility. The goals of normalization require that both precision and accuracy be examined. No single metric captures either of these attributes perfectly and they are likely to be region, study and population dependent. Still, meaningful improvements in both precision and accuracy can be achieved through careful construction of processing pipelines, as with the multi-tool T1EPI pipeline developed at Maryland, which consistently shows the best overall performance. Although differences between the three SPM-based pipelines were smaller and more variable, the widely used T1EPI_standard_ pipeline was the most precise of the three, particularly with cerebral voxels, whereas the recently developed EPIO_calhoun_ pipeline was the least accurate. These three pipelines can be readily improved (e.g., by developing improved EPI templates or incorporating fieldmap correction) and we encourage researchers to invest the resources necessary to do so. We anticipate that the performance metrics described here will prove useful for validating pipelines developed in the future.

## CONTRIBUTIONS

J.F.S. envisioned and designed the study. J.F.S. developed and implemented the image registration/normalization pipeline. C.M.K. and J.F.S. collected data. J.F.S. processed data. J.F.S. and J.H. analyzed data. J.F.S. and A.J.S. interpreted data. J.F.S and A.J.S. wrote the paper. J.F.S. created figures and tables. A.J.S. funded and supervised all aspects of the study. All authors contributed to reviewing and revising the paper and approved the final version.

## ACKNOWLEDGEMENTS

Authors acknowledge guidance on pipeline parameters from V. Calhoun and assistance from K. DeYoung, L. Friedman, G. Kim, and S. Padmala. This work was supported by the University of Maryland, College Park and National Institutes of Health (DA040717 and MH107444). Authors declare no conflicts of interest.

## DATA AVAILABILITY

Select data have been or will be publicly shared using NeuroVault.

## Footnotes

1 We use the terms ‘atlas’ to indicate a standard stereotaxic space and ‘template’ to indicate a reference image within that space. For example, the MNI-ICBM 2009 atlas consists of T1-weighted, T2-weighted, and proton density-weighted templates of varying resolutions (0.5-1 mm^3^) in a common coordinate system(http://www.bic.mni.mcgill.ca/ServicesAtlases/ICBM152NLin2009).

2 Similar techniques have been applied to positron emission tomography (PET) data.

3 We note that while the SPM segmentation used in creating the brain mask could also be used here for convenience, BBR coregistration was validated with FAST.

4 **Using participant 34 as the reference**: On average, images from the T1EPI_Maryland_ pipeline were more strongly correlated with the reference than those from the EPIOCalhoun and T1EPI_Standard_ pipelines (*ts*(47) > 2.04, *ps* < .05), but not those from the EPIO_Standard pipeline_ (*p* = .10). Images from the two EPIO pipelines were more strongly correlated than those from the T1EPI_Standard_ pipeline (*ts*(47) > 2.24, *ps* < 0.03), but the difference between them was not significant (*p* = .71). Using participant 16 as the reference: Images from the T1EPI_Maryland_ pipeline were more strongly correlated than those from the other three pipelines (*ts*(47) > 5.48, *ps* < .001. Other differences were similar to those when using participant 34 as the reference image.

5 **Using participant 34 as the reference**: On average, images from the two T1EPI pipelines were more strongly correlated with the reference than those from the two EPIO pipelines (*ts*(47) > 5.41, *ps* < .001), but did not significantly differ from one another (*p* = .77). Images from the EPIO_Standard_ pipeline outperformed those from the EPIO_calhoun_ pipeline (*t*(47) = 7.37, *p* < .001). **Using participant 16 as the reference**: The overall pattern of results was identical, but here the T1EPI_Matyiand_ pipeline outperformed the T1EPI_standard_ pipeline (*t*(47) = 4.95, *p* < .001.

6 All peaks fall within the areas described except [54,60,8], the LOC peak from the EPIOStandard data. Though ~4mm away from the other peaks, this coordinate is identified as “Middle Temporal Gyrus, temporoccipital” part by the atlas rather than LOC.

7 Likewise, within the right prefrontal cluster, data normalized using the T1EPI_Maryland_ pipeline yielded the strongest effects at one local maximum (*x* = 51, *y* = 12, *z* = 21; *N*_*eff*_ increase of 41.6-49.3%) and the weakest effects at the other (*x* = 53,*y* = 20, *z* = 25; *N*_*eff*_ decrease of 16.5-28.0%).

8 Indeed, the whole volume precision of EPIO_calhoun_ seems due to its superior performance in *extra*cerebral regions (i.e., outside of the eroded mask) where its average participant correlation to the grand average (EPIO_calhoun_ *r* = 0.776) was significantly larger than T1EPI_Standard_ (T1EPI_Standard_ *r* = 0.758, *t*(48) = 5.20, *p* < 0.001) as well as T1EPI_Maryland_ (T1EPI_Maryland_ *r* = 0.746, *t*(48) = 4.83,*p* < 0.001).

